# Bypass of DNA Interstrand crosslinks by a Rev1-DNA polymerase ζ complex

**DOI:** 10.1101/2020.05.09.085902

**Authors:** Rachel Bezalel-Buch, Young K. Cheun, Upasana Roy, Orlando D. Schärer, Peter M. Burgers

## Abstract

DNA polymerase ζ (Pol ζ) and Rev1 are essential for the repair of DNA interstrand crosslink (ICL) damage. We have used yeast DNA polymerases η, ζ, and Rev1 to study translesion synthesis (TLS) past a nitrogen mustard-based ICL with an 8-atom linker between the crosslinked bases. The Rev1-Pol ζ complex was most efficient in complete bypass synthesis, by 2-3 fold, compared to Pol ζ alone or Pol η. Rev1 protein, but not its catalytic activity, was required for efficient TLS. A dCMP residue was faithfully inserted across the ICL-G by Pol η, Pol ζ, and Rev1-Pol ζ. Rev1-Pol ζ, and particularly Pol ζ alone showed a tendency to stall before the ICL, whereas Pol η stalled just after insertion across the ICL. The stalling of Pol η directly past the ICL is attributed to its autoinhibitory activity, caused by elongation of the short ICL-unhooked oligonucleotide (a six-mer in our study) by Pol η providing a barrier to further elongation of the correct primer. No stalling by Rev1-Pol ζ directly past the ICL was observed, suggesting that the proposed function of Pol ζ as an extender DNA polymerase is also required for ICL repair.

## Introduction

DNA Polymerase ζ (Pol ζ) is a multi-subunit B-family DNA polymerase enzyme that is involved in translesion DNA synthesis (TLS) (1). Unlike the other B family members (Pol α, Pol δ, Pol ε), which show a high replication fidelity, Pol ζ has a low fidelity (2,3). Pol ζ is a four-subunit DNA polymerase (Rev3, Rev7, Pol31, Pol32), sharing the Pol31 and Pol32 subunits with Pol δ (4,5). Its interaction with the replication clamp PCNA increases its activity, presumably by increasing the processivity of the enzyme (5,6). Rev1 DNA polymerase is a Y family DNA polymerase with a unique dCMP transferase activity (7). However, while *Saccharomyces cerevisiae REV1* is absolutely required for damage-induced mutagenesis (8), its catalytic activity is not (9), suggesting that the non-catalytic functions of Rev1 are important. Therefore, Rev1 is considered mainly as a scaffold protein onto which the mutasome is assembled. It promotes mutagenesis by essential interactions with other factors, including mono-ubiquitinated PCNA and Pol ζ (10-13).

Cells deficient for Pol ζ or Rev1 are sensitive to DNA damage and are among the most hypersensitive to treatment with interstrand cross-linking (ICL) agents, suggesting that they have a key role in ICL repair (14-19). DNA ICLs covalently link two strands of the double helix, thereby providing a complete block to both the DNA replication and the transcription machineries. ICLs can be formed by endogenous sources as well by antitumor agents such as nitrogen mustard and cisplatin (20,21). The removal of ICLs from genomes necessarily requires a complex repair process and several pathways for ICL repair have been described. Although ICL repair also occurs in the G1 phase of the cell cycle, the removal of ICLs is most critical during S-phase, where they provide an absolute block to replication (22). ICLs can be encountered by the collision of two replication forks (23), or a unidirectional fork can traverse the ICL in a FancM-dependent step followed by priming and fork resumption downstream of the ICL (24). Regardless, all ICL repair pathways require an unhooking step to separate the two crosslinked strands and a DNA synthesis step to restore the duplex. Studies of the repair of site-specific plasmid-based ICLs in *Xenopus* egg extracts have provided a biochemical framework for understanding ICL repair and illustrate what roles DNA polymerases play in ICL repair (23). Activation of the Fanconi anemia (FA) pathway at the ICL leads to the recruitment of the ERCC1-XPF and possibly other endonucleases to unhook the crosslink from one of the two strands (25,26). The unhooked ICL may be trimmed further by an exonuclease such SNM1A/Pso2 (27). The unhooked and processed ICL is then bypassed by translesion synthesis (TLS) DNA polymerases to generate one continuous duplex that can be used as a template to fix the second strand (23). While this FA-dependent pathway can act on most ICLs, several variants of this general pathway exist for ICLs formed by psoralen, abasic sites or acetaldehydes (28,29).

Studies in the xenopus system showed that depletion of Rev1 from the extract did not affect the efficiency and kinetics of insertion at the ICL site, however, it did have an inhibitory effect on the extension reaction (13). Similarly, depletion of the Rev7 inhibited the extension of the repair product past the insertion opposite a cisplatin ICL, in agreement with the known role of Pol ζ as an extender polymerase (23). Biochemical studies have shown that various TLS polymerases can bypass ICLs (21,30-32), but conclusive studies of the bypass activity of purified Pol ζ and Rev1, are missing. In this paper, we provide critical new information regarding the functions of Pol ζ and Rev1 on an ICL substrate that mimics an unhooked nitrogen mustard lesion (30,33). Our studies show that Rev1 stimulates ICL bypass by Pol ζ, but its enzymatic activity is largely dispensable. Both Pol η and Rev1-Pol ζ, but not Pol δ can mediate ICL bypass. However, Pol η shows very strong stalling directly past insertion at the ICL. In contrast, Rev1-Pol ζ proceeds more efficiently past the ICL, suggesting that the enzyme is suitable for complete TLS.

## Results

### Generation of ICL substrates for PCNA-mediated polymerase bypass

We and others have previously shown that the structure of the ICL that links two bases as well as the length of the duplex around an ICL affects bypass by DNA polymerases (30,31,33). We used our previously published method to generate a stable nitrogen mustard ICL mimic with an 8 atom crosslink through a double reductive amination reaction of an aldehyde ICL precursor with dimethylethylenediamine embedded in a 6mer duplex (8a-6bp ICL, Figure 1A) (34,35). To the initially obtained ICL on a 39mer template, we ligated 5’ and 3’ biotinylated oligos to generate a 93mer substrates with biotinylated ends. Our design was based on the consideration that i) when replication forks stall, about 20 nucleotides remain unreplicated between the leading strand 3’-end and the ICL, and ii) that nucleolytic processing of the non-template strand leaves the unhooked ICL in a duplex of only a few nucleotides (23). The biotin moieties at either template terminus were introduced to bind streptavidin to form blocks to prevent PCNA from sliding off the DNA (6). The control substrate (undamaged DNA) lacks the ICL and the 6 nucleotides of dsDNA, which would not form a stable duplex with the template strand in the absence of the ICL.

**Figure 1.**
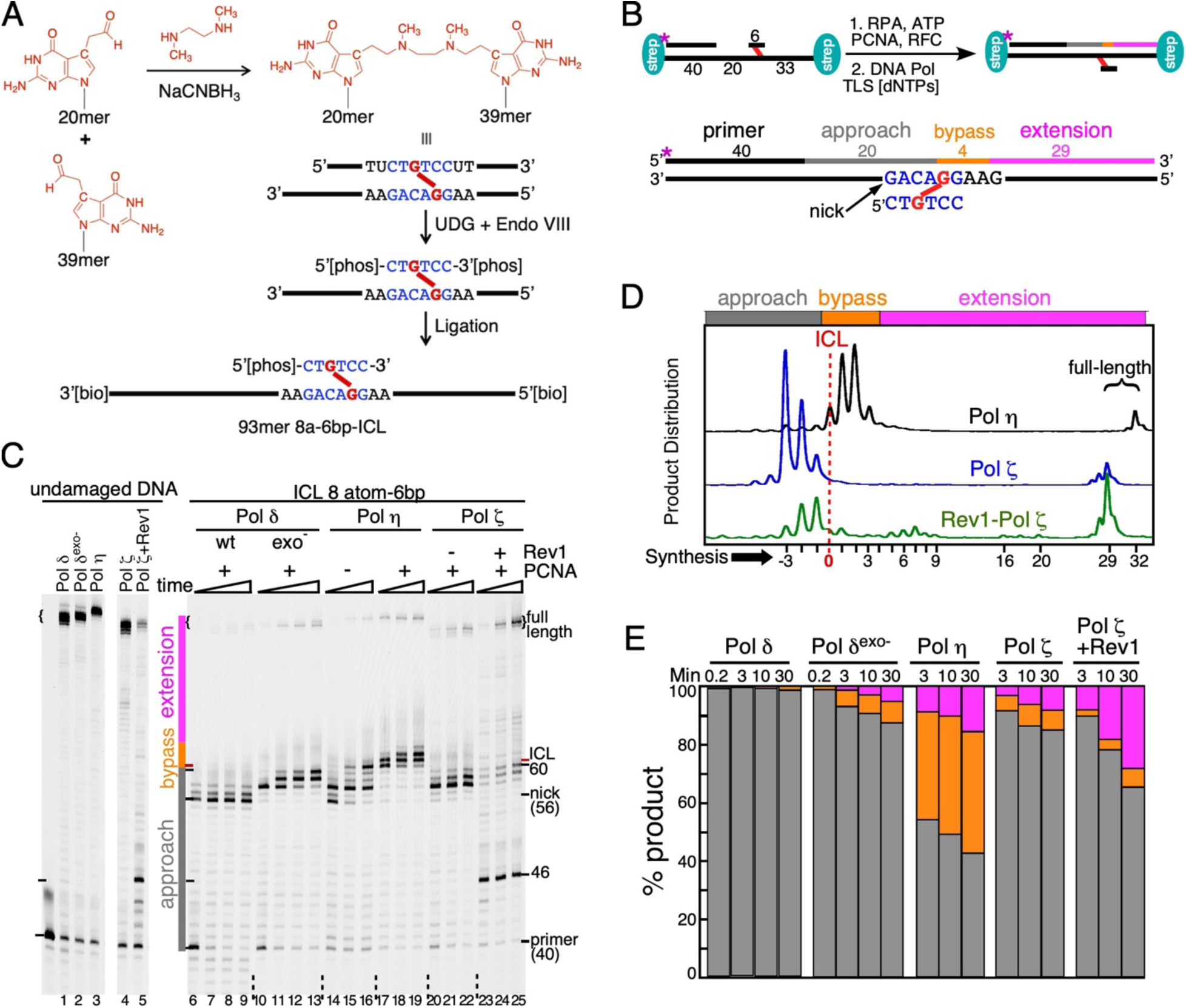
Translesion DNA synthesis of an ICL by yeast DNA polymerases. (**A**) Synthetic scheme for the 93-ICL+6bp Substrate. UDG, uracil-DNA-glycosylase; Endo VIII, *E. coli* endonuclease VIII. See Materials and Methods for details. **(B)** Top, schematic overview of the DNA substrate and the assay set-up. Bottom, designation of specific classes of elongation products: Approach, gray; Bypass, orange; Extension, pink. TLS [dNTPs] are concentrations that prevail under damage response conditions in yeast. See Materials and Methods for details. **(C)** Time course of replication of control DNA (lanes 1-5) or ICL DNA by the indicated DNA polymerase, with or without PCNA for Pol η, and with or without Rev1 for Pol ζ. The gel was cut up for easier visualization (the entire gel is shown in Supplementary Figure S2A). **(D)** The extension products, from a 53-mer size up to full-length, obtained by scanning lanes 19, 22 and 25, were plotted. The presence of the streptavidin block at the template 5’-end limits full replication by the bulky Pol ζ and Rev1-Pol ζ enzymes, compared to Pol η. **(E)** Quantification of the replication products from (C) into three classes.

### In vitro bypass of an ICL by Pol η and Rev1-Pol ζ

We have previously shown that Pol ζ can be purified as a stable 5-subunit complex together with Rev1 (36). A complex with the identical activity can also be obtained by simply mixing Pol ζ together with Rev1, thereby reconstituting the 5-subunit complex. In this paper, we have reconstituted the various complexes of Rev1-Pol ζ by mixing Rev1, or Rev1 mutants, together with wild-type four-subunit Pol ζ.

The DNA substrates were pre-incubated with the single-stranded DNA binding protein RPA (replication protein A), and PCNA (proliferating cell nuclear antigen) was loaded by RFC (replication factor C) and ATP, as shown in Figure 1B. The reaction was started by addition of the individual DNA polymerases. dNTPs were present at concentrations that prevail during the DNA damage response in yeast (36). Replication of the undamaged control DNA was carried out very efficiently, with the exception of replication by Rev1-Pol ζ (lane 5, discussed below). The presence of the streptavidin moiety at the template 5’-end sterically limits full extension by Pol δ (lanes 1,2) and Pol ζ (lanes 4,5), compared to Pol η (lane 3), and this pattern was also displayed with the ICL-containing substrates.

Pol δ stalled mainly at the nick position, four nucleotides prior to encountering the crosslinked G residue (Figure 1C, lanes 6-9), consistent with its known idling function at DNA nicks (37). However, about 2-3% full-length product was observed at every time point, which could be due to incomplete crosslinking of the substrate or a low level of bypass by Pol δ (see also Supplementary Figure S2A). This non-specific background observed by wild-type Pol δ was subtracted from the percentage of extension products observed with the TLS enzymes. First, we eliminated the proofreading function of Pol δ (Pol δ-exo^-^, D520V). This reduced idling at the nick (37), but still allowed only minimal bypass synthesis over the 30 min time-course of the assay (Figure 1C, lanes 10-13; Figure 1E).

Based on previous studies of TLS by Pol η, we have divided the replication products into three classes for the purpose of quantification (33). Products up to the ICL are designated as “**approach**”, those that have inserted a nucleotide opposite the crosslinked G, plus an additional three nucleotides past the ICL are designated as “**bypass**”, and those longer than that up to full-length are designated as “**extension**”. Quantification is shown in Figure 1E. The bypass of this particular ICL has been studied previously, however these model studies were carried out with Pol η alone, without the PCNA clamp and at low salt concentrations (33). TLS by Pol η alone was most efficient at low salt (50 mM NaCl, Supplementary Figure S2B). For sake of consistency, all studies in this paper were carried out at 100 mM NaCl. Under these conditions, the limited bypass of the ICL by Pol η was stimulated substantially by PCNA (Figure 1C, compare lanes 14-16 with 17-19 & Figure S2B, compare lanes 7-9 with 10-12, and 13-15 with 16-18). However, as observed previously (33), the majority of products are still stalled at the +1 to +3 positions. In contrast, Pol ζ stalled predominantly prior to the ICL, similar to proofreading-defective Pol δ (lanes 20-22). However, significant bypass and extension was observed. Notably, the lack of products at the +1 to +3 positions, together with the presence of fully replicated DNA indicates that, once a nucleotide has been inserted opposite the ICL-G position, this product is favorably accommodated in the Pol ζ binding site to allow continued extension. This remarkable difference between Pol η and Pol ζ is shown graphically in Figure 1D.

Three dramatic changes were observed with Rev1-Pol ζ as TLS enzyme compared to Pol ζ alone. First, strong pause sites were observed during replication of normal (undamaged) DNA (Figure 1C, lanes 4 and 5). The inhibition of Pol ζ activity by Rev1 on undamaged DNA is the subject of a different study (Bezalel-Buch, R. and Burgers, P. unpublished data). Replication of the ICL-DNA also shows these ICL-independent distant pause sites, in addition to ICL-dependent pause sites directly ahead of the crosslink. Second, synthesis past the ICL is enhanced by the inclusion of Rev1 (Figure 1C, lanes 23-25). Third, once bypass has been achieved, further replication of the downstream template is once more inhibited as is evident from the presence of multiple, distant stall sites. After 30 min, the total extension products by Rev1-Pol ζ are about 30% compared to 10% with Pol ζ alone (Figure 1C-E).

### The catalytically inactive form of Rev1 stimulates Pol ζ-mediated ICL bypass

The data in Figure 1 show clearly that Rev1 has an important role in Pol ζ-mediated ICL bypass and extension. To assess whether this is due to the catalytic function of Rev1 or to its proposed scaffolding function, we repeated the TLS assay with Rev1^cd^ (Rev1-DE467,468AA), a catalytic-inactive mutant of Rev1 (Figure 2A, (9)). Remarkably, the efficiency of ICL bypass and the extension by Rev1^cd^-Pol ζ was higher than by Pol ζ alone, and it is diminished only slightly from that observed with wild-type Rev1-Pol ζ (Figure 2B). The major difference is the stronger accumulation of stalled products directly prior to the ICL, suggesting the participation of the catalytic activity of Rev1 at the ICL. Interestingly, the major lesion-independent stall site at position 46 is still present albeit reduced by 30-55% (Figure 2A, lane 3 *versus* 4, and lanes 11-13 *versus* 14-16). In addition, post-ICL stall sites are somewhat reduced, including a minor site at position 70. In order to determine the possible participation of the catalytic activity of Rev1, we sequenced the fully replicated products from a 60 min assay, which resulted in >60% full-length product accumulation (not shown) (Figure 2C, Supplementary Figure S3). Remarkably, while bypass of the major stall site at position 46 (across a dC template) is not associated with dCMP misincorporation (Supplementary Figure S3), bypass of the minor stall site at position 70 (across a dC template) is associated with misincorporation of dCMP with a 15-20% efficiency (3 determinations). Misincorporation is carried out by Rev1’s dCMP transferase activity, since it is not present with Pol ζ alone, and eliminated with Rev1^cd^-Pol ζ (Figure 2C). Therefore, the inhibitory mechanisms of Rev1 on non-damaged DNA involve both its catalytic and scaffolding functions. Within the sensitivity of detection (>95%), all three enzymes (Pol ζ, Rev1-Pol ζ, Rev1^cd^-Pol ζ) faithfully introduced a dCMP across the ICL-G position (Figure 2C).

**Figure 2.**
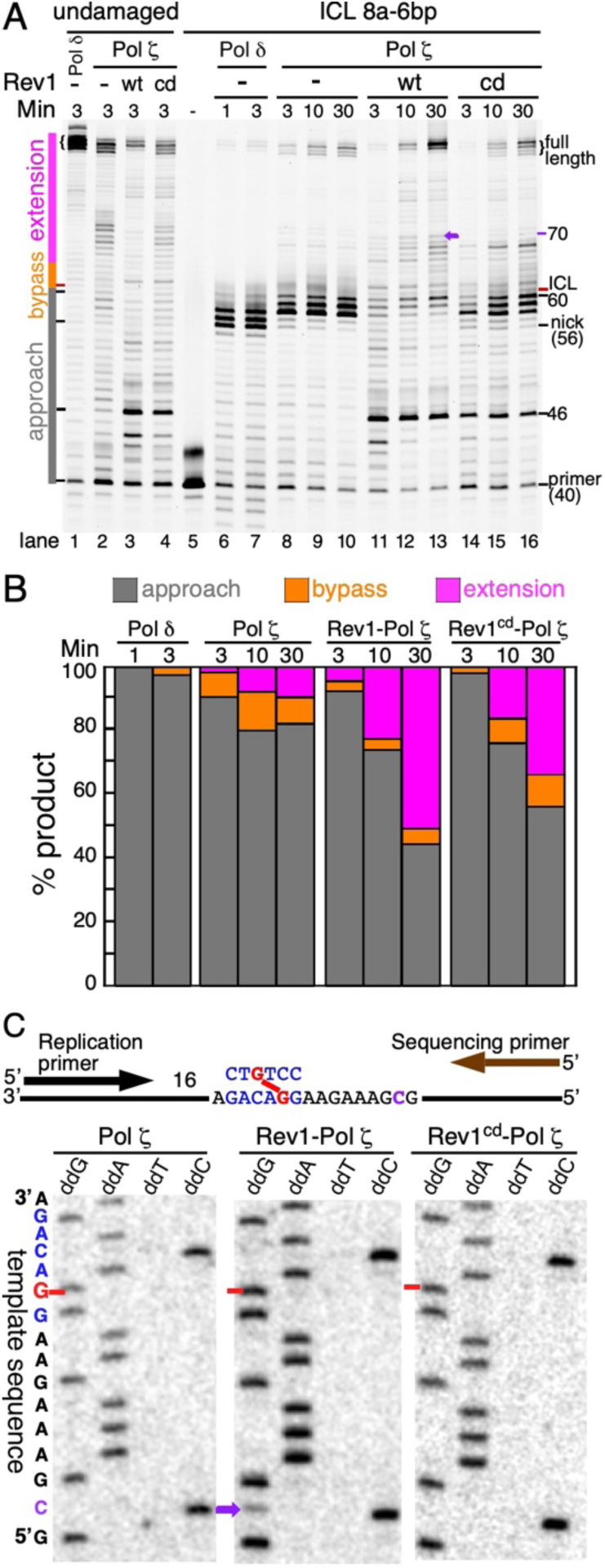
Catalytically inactive Rev1 stimulates ICL TLS by Pol ζ. **(A)** Time course of replication of control DNA (lanes 1-4) or ICL DNA by the indicated DNA polymerase, with or without Rev1 for Pol ζ. Rev1^cd^ is catalytic-dead (D467A/E468A). **(B)** Quantification of the replication products into three classes. **(C)** ICL bypass product sequence analysis. Top, schematic map of the assay. Full-length replication products were isolated from 60 min assays. Bottom, sequencing analysis. The red arrow indicates the ICL-dG template position, which is replicated by insertion of dCMP, which is sequenced as a dG. The purple arrow, also shown in (A) at position 60, indicates a position where Rev1, but not Rev1^cd^ misincorporates dCMP across from template dC with a frequency of 15-20%.

### Distance of the primer terminus from the nick position influences TLS by Rev1-Pol ζ

It is not entirely clear where and when during DNA replication and translesion synthesis, a TLS polymerase takes over DNA synthesis. Is it when replication stalls ∼20 nt before the ICL (on the leading strand), or only when the ICL position is reached? A recent study in *Xenopus* egg extract suggests that the approach is carried out by a replicative DNA polymerase, up to one nucleotide before the ICL position, and bypass and extension is performed by a complex of Rev1 and Pol ζ (13). We determined the efficiency of ICL bypass as a function of the distance between the primer terminus and the ICL (Figure 3). The three primers were 40 nt in length and were positioned with their 3’-termini at either 20 nt, 8 nt, or 6 nt from the ICL position. Pol η showed efficient bypass with all three substrates, and stalling at the +1 to +3 positions past the nick remained unaffected. Bypass by Pol ζ alone was stimulated slightly when the primer was positioned closer to the ICL (compare lanes 6, 12, and 18). However, we noticed a dramatic improvement in bypass synthesis by Rev1-Pol ζ when the primer terminus positioned was closer to the ICL (compare lanes 7, 13, and 19). To a large extent this increase in efficiency can be attributed to the elimination of the ICL-independent strong stall sites by hybridization of the closer primers. As expected, once the ICL was bypassed, Rev1-Pol ζ stalled similarly at downstream template positions. The distance between the primer terminus and the ICL did not affect the identity of the nucleotide incorporated opposite the ICL-G. With both the −20 and the - 6 primer (with relation to the ICL), this was a dCMP residue, introduced either by Pol η, by Pol ζ, or by Rev1-Pol ζ (Figure S3 & S4).

**Figure 3.**
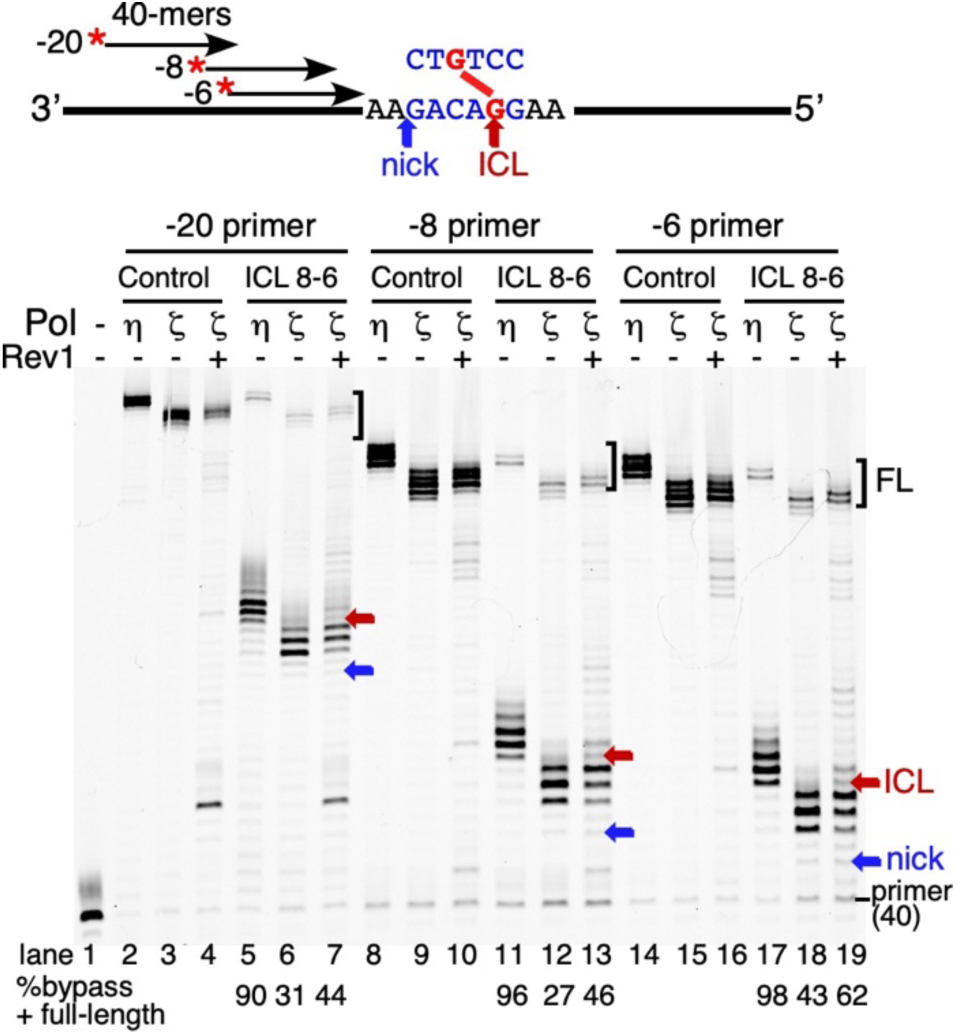
Primer distance from the ICL influences TLS efficiency. Top, schematic map of the three different 40-mer primers, the nick location and the ICL position. Bottom, gel analysis of the assays, with either the control or ICL substrates. Replication was carried out for 15 min with either Pol η, Pol ζ, or Rev1-Pol ζ, as indicated. Blue and red arrows indicate nick and ICL positions, respectively, for the different substrates. At bottom, quantification of the % bypass + % extension products (See Figure 1B).

### The interplay between different DNA polymerases in ICL bypass

While it is well recognized that Pol ζ and Rev1 are indispensable for the TLS of ICLs in eukaryotic cells, Pol η can contribute to the efficiency of this process (38). Switching between Pol η and other DNA polymerases has been observed with pyrimidine dimer lesions (39-41). Does switching also occur during the TLS of ICLs, and with what efficiency? This is of particular interest since Pol η stalls at positions past the ICL, which in theory should provide good substrates for extension by Pol ζ or Rev1-Pol ζ. Therefore, we investigated a potential collaboration between DNA polymerases in ICL bypass. In the assay we used all combinations of Pol δ, Pol η, Pol ζ and Rev1 (Figure 4A).

**Figure 4.**
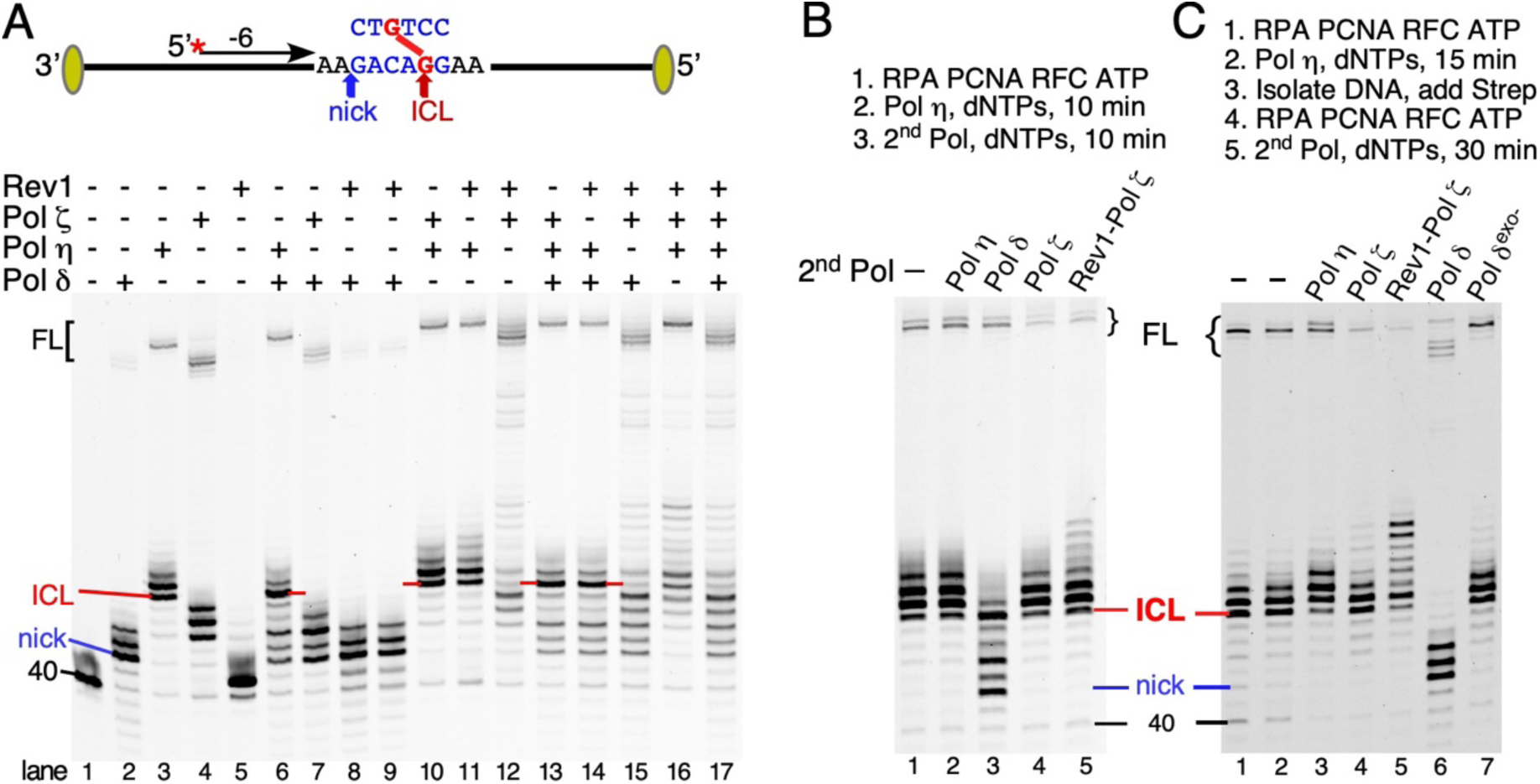
Polymerase exchange analysis in ICL bypass synthesis. The ICL substrate with the −6 primer was used. **(A)** Reactions were carried out for 15 min with either the individual DNA polymerases, or with a combination, which was added as a pre-formed mixture, as shown. Note that all lanes containing Pol δ show degradation products shorter than the 40 mer primer. These products were less than 10% of total. **(B)** Two-stage reaction with Pol η in the first 10 min incubation, followed by the indicated DNA polymerase in the second 10 min incubation. Lane 1, no second incubation; lane 2, incubation with Pol η continued. **(C)** Two stage ICL bypass assay. After the first incubation, the DNA was isolated (materials and methods), followed by second stage assay with the indicated DNA polymerase. Lane 1, product after first stage; lane 2, second stage incubation without added polymerase.

Lanes 2-5 represent the reactions with one DNA polymerase. These data recapitulate the data shown in Fig. 1, with Pol δ stalling at the nick, Pol η at the 0 (ICL), +1 and +2 positions, Pol ζ 1-3 nucleotides before the ICL, and Rev1 alone stalling after incorporation of one single nucleotide. With the exception of Rev1-Pol ζ, the combination of two DNA polymerases did not increase bypass and extension significantly. In fact, the presence of Pol δ inhibited bypass by Pol η and Pol ζ (compare lanes 6, 7 with 3,4). Surprisingly, Pol η plus Pol ζ did not yield a substantial increase in extension products compared to Pol η alone (compare lane 10 with 3).

As shown earlier, only the Rev1-Pol ζ complex showed an increase in bypass and extension (lane 12). Next, three DNA polymerases were tested. Again, Pol δ showed only an inhibitory activity on bypass (lanes 13-15), by degrading extension products through its 3’-exonuclease activity as shown in Figure 1C. Surprisingly, like observed with the Pol η-Pol ζ couple, the combination of Pol η and Rev1-Pol ζ yielded only a marginal increase in bypass compared to Rev1-Pol ζ alone (compare lanes 16 and 12). Significantly, most products are stalled at the ICL and 1-2 nucleotides past the ICL, which is evidence of Pol η activity (lane 3). Finally, the four-polymerase bypass reaction again showed the inhibitory effect of Pol δ on TLS (lanes 16, 17).

We were surprised by the observation that the combination of Pol η and Rev1-Pol ζ did not give the anticipated synergy in bypass. We had predicted that this combination would give excellent TLS because Pol η alone would produce mostly products that stalled past the ICL, while Pol ζ or Rev1-Pol ζ showed no stalling at these positions. Therefore, we expected that Pol ζ or Rev1-Pol ζ would readily extend the Pol η intermediates to full-length products (Figure 1D). However, this was not observed. We carried out a two-step bypass assay, in which the ICL product was replicated by Pol η in the first stage, and then further extended with other DNA polymerases in the second stage (Figure 4B). While Pol δ was inhibitory in the second stage, degrading the bypass products by Pol η (lane 3), Rev1-Pol ζ showed only minimal extension of the bypass products (lane 5).

We next examined the possibility that the Pol η was bound up in a stalled, non-exchangeable (frozen) complex with the +1 to +3 bypass products, blocking access by Rev1-Pol ζ. This hypothesis needed testing, in spite of the observation that the 3’-exonuclease activity of Pol δ did have access to these stalled products (Figure 4B, lane 3). After the first stage, the DNA was reisolated by ethanol precipitation and subjected to the second stage assay (Figure 4C). Analogous results to those in Figure 4B were obtained. While Rev1-Pol ζ promoted significant propagation of the Pol η bypass products, it was still limited, yielding very few full-length products during the 30 minute incubation.

Translesion synthesis in the cell requires mono-ubiquitination of PCNA at Lys-164 (42). Mono-ubiquitinated PCNA shows increased binding to Rev1 and it stimulates Rev1 activity in certain sequence contexts (10,12,43). However, mono-ubiquitinated PCNA does not stimulate Pol ζ activity (10). The main proposed function of monubiquitination is to recruit Rev1, and thereby Pol ζ to damaged chromatin for TLS. In a control experiment, in which PCNA and ubiquitinated PCNA were compared as accessory factors for TLS of the ICL, we detected no significant difference in bypass efficiency (Supplementary Figure S5).

### Autoinhibitory activity of Pol η during ICL bypass

We next investigated whether the bypass products made by Pol η were mere elongation products of the 40-mer primer, or whether other changes to the ICL substrate had occurred that were responsible to led to an inhibition of the formation of full-length products. If primer elongation, for instance to the +2 position, were the only result of replicating the ICL substrate with Pol η, then that +2 product should be equivalent in activity to a substrate made by hybridizing the +2 primer to the ICL template. In the experiment in Figure 5, different length primers, with the same 5’-end but with their 3’-termini at the −6 position (the standard primer), at 0 (across the ICL), or at +2 or +6 position, were hybridized to the ICL template. Their reactivities were assessed with each of the four enzymes. For the purpose of this experiment, we defined replication product from ten nucleotides past the ICL to full-length as extension products. When the series of substrates was tested with Pol η, we saw some improvement in full-length product with the +2 primer compared to the standard −6 primer (Figure 5A), but stalling was still a major problem. Only the +6 primer escaped the problem of stalling. Pol δ extended only the +6 primer efficiently, suggesting that the double-stranded DNA binding cleft of Pol δ is very sensitive to the ICL-induced structural alterations from Watson-Crick base-pairs (Figure 5B). Nevertheless, even the +6 substrate was compromised for extension by Pol δ. About 20 % was subject to degradation by Pol δ’s proofreading activity, as follows from the generation of products at positions around the nick. This was observed only with the ICL substrate and not with the control substrate (compare lanes 14 and 15).

**Figure 5.**
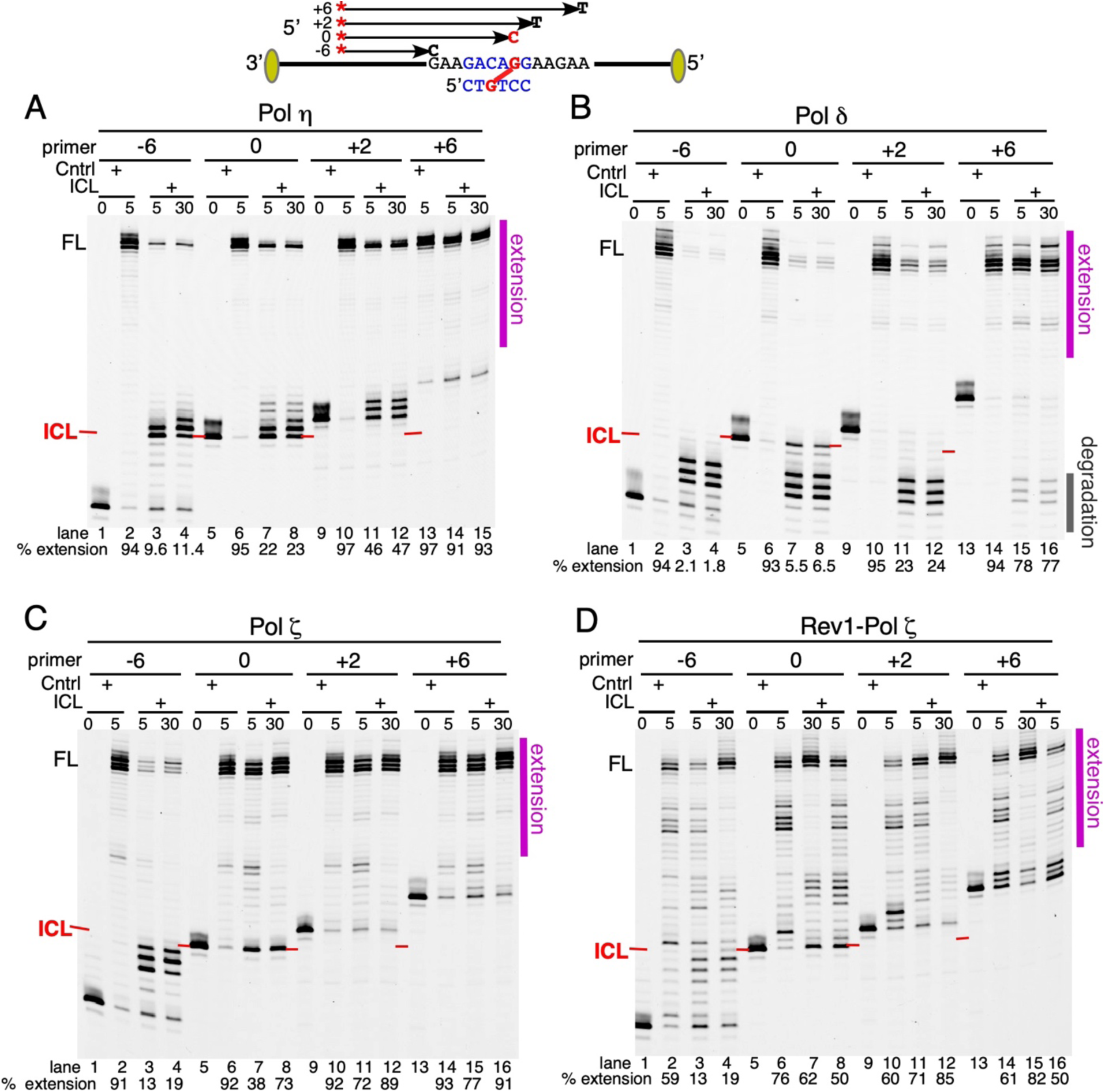
Extension of primers hybridized beyond the ICL. The standard primer (−6) and primers at the ICL position (0) and beyond (+2, +6), were used as substrates for extension by Pol η **(A)**, Pol δ **(B)**, Pol ζ **(C)**, and Rev1-Pol ζ **(D)**. The extension products from 10 nucleotides beyond the ICL (or the corresponding position on the control template) up to full-length, were defined as extension products and given under each panel. In panel (B), the main degradation products by Pol δ are indicated to the right. The FAM-primers show variable ghosting of fluorescence at positions ∼1-2 nt higher (compare the identical substrate in panel A, lane 5 with that in panel C, lane5). This ghosting phenomenon is only for unreplicated primers and disappears upon replication.

In sharp contrast, Pol ζ extended even the (0) substrate with very high efficiency, while the −6 substrate shows the usual stalling just prior to the ICL (Figure 5C). Rev1-Pol ζ also fully extended the primer opposite the ICL as well as the +2 and +6 primer (Figure 5D). The continued stalling further downstream of the ICL is a result of the negative regulatory activity of Rev1 on Pol ζ, and was also observed with the control template (Figure 5D, e.g. compare lane 10 with 11).

One likely cause for the autoinhibitory activity of Pol η during ICL bypass could be that the 6-mer, which is crosslinked to the template is extended by Pol η. Consequently, after ICL bypass, further elongation would require strand displacement synthesis through the extended 6-mer strand, which would be inhibitory for these TLS polymerases (30). The assay described in Figure 6 lends support to this hypothesis. We incubated the unprimed ICL substrate, or the ss93-mer as control, with Pol η, followed by hybridization of the standard primer (−6), and then a second incubation with Pol η. The FAM-primer and its extension products were detected by FAM fluorescence (Figure 6A), while all DNAs were detected with SYBR-gold staining (Figure 6B). The appearance of a Pol η-dependent slow-migrating species in (B) with the unprimed 93-ICL+6bp, but not with the un-crosslinked 93-mer template, is evidence that indeed, Pol η used the covalent 6-mer as a primer for extension, yielding a 93-ICL+37bp product upon replication to the end of the available template (Figure 6B, lanes 7,8). We propose that this extension product is inhibitory for full extension of the FAM-primer not only by Pol η, and also by Pol ζ or Rev1-Pol ζ.

**Figure 6.**
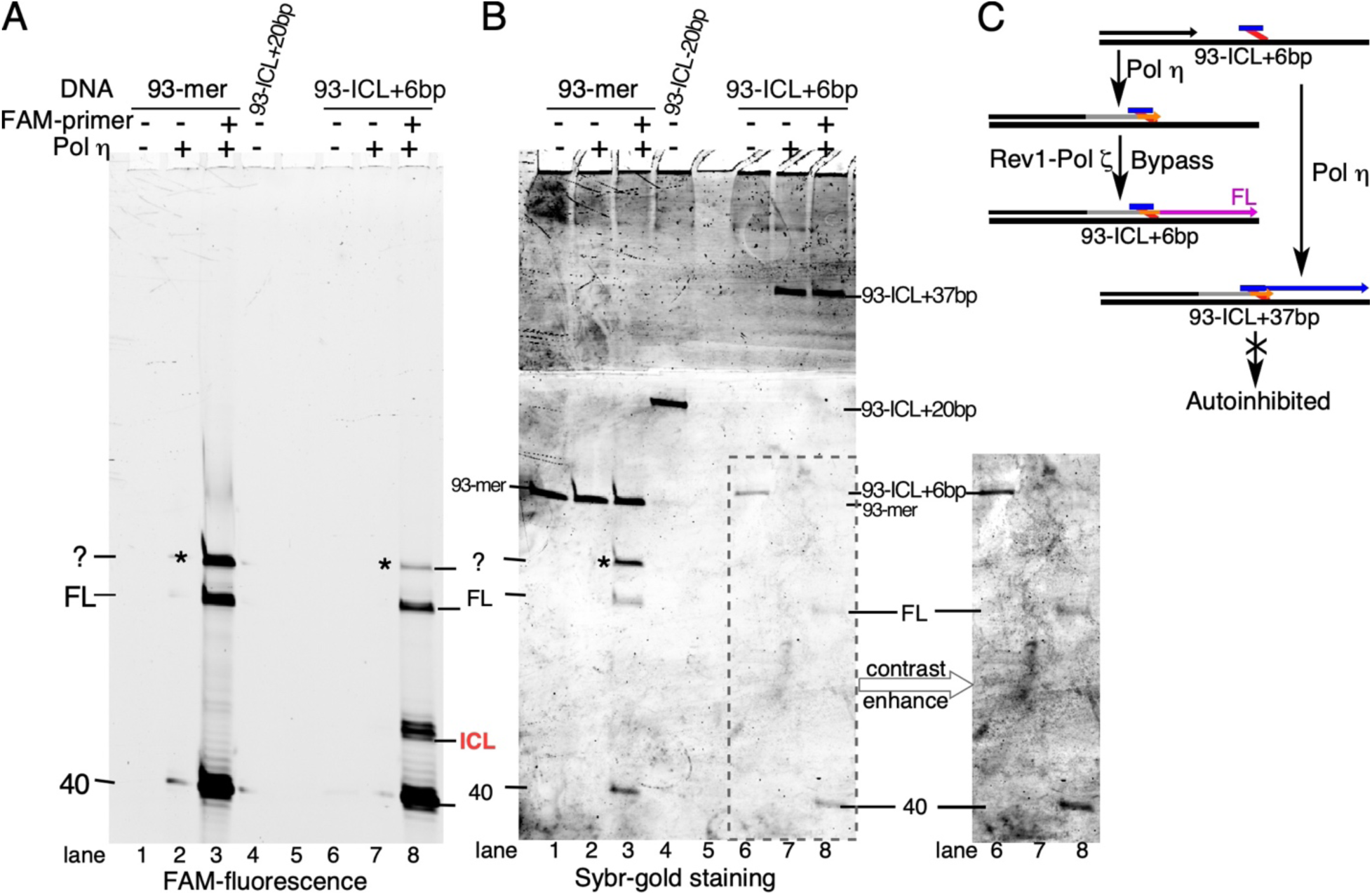
Extension of the crosslinked six-mer by Pol η. Standard assays were increased to 20 μl with DNA concentrations increased to 20 nM, and the proteins accordingly. In the first incubation, assays were carried out with the indicated unprimed template. After 30 min with Pol η, reactions in lanes 3 and 8 were stopped for 1 min at 70 °C, and the (−6) 5’-FAM-40-mer primer was hybridized (a 10-fold excess of primer was used). Assays were reinitiated with RPA, PCNA, RFC and Pol η for 30 min, and analyzed on a 12% PAGE-urea gel, containing 25% formamide. The gel was first scanned in the Typhoon for FAM fluorescence **(A)**, and subsequently stained with SYBR GOLD, after the gel was cut in two pieces. It was then scanned in the Typhoon, using the EtBr setting **(B)**. In the inset at right, the bottom of the SYBR GOLD-stained gel was contrast enhanced to show weak FL synthesis in lane 8. 93-ICL+37 is the expected size if the covalently attached 6-mer performs as a primer for extension by Pol η. 93-ICL+20bp (lane 4) is used as a size marker (Figure 1A). *These replication products are a consequence of the high primer/template ratio of 10 in this experiment. They were not observed when primer and template were equimolar.

## Discussion

For over a decade, a consensus model has emerged for TLS in eukaryotic cells (44-46). This process is usually carried out by the tandem action of two DNA polymerases. The first DNA polymerase, the inserter, incorporates one or more nucleotides opposite the lesion, whereas the second DNA polymerase, the extender, continues replication. Depending on the type lesion, the inserter DNA polymerase is usually a Y-family DNA polymerase, such as Pol η, Pol κ, or Pol ι in mammalian cells, whereas Pol ζ is generally considered to be an extender DNA polymerase (1). In yeast, Pol η is the only classical “inserter” Y-family enzyme. The second Y-family enzyme, Rev1 has a specialized obligatory function with Pol ζ, and it does so without generally requiring its unique deoxycytidyl transferase activity, including during TLS of cis-platin induced DNA adducts (47,48). One caveat to these genetic observations with Rev1 is that they do not distinguish between *intra-* and *inter*strand DNA lesions caused by cis-platin.

In our study of a highly replication-blocking ICL, the role of Pol ζ as an extender is readily apparent. Once the insertion of a nucleotide across the ICL has been made, extension by Pol ζ was facile, regardless of the presence of Rev1 (Figure 5C, D). But the question as to which DNA polymerase is involved in the insertion process remains. Based on Rev1 and Pol ζ depletion studies in *Xenopus* egg extracts, it has been suggested that Rev1-Pol ζ is not responsible for insertion across the ICL, but it is for the extension step (13,23). Therefore, it follows that another DNA polymerase should be responsible for the insertion step and Rev1-Pol ζ for extension.

The purpose of the current study was two-fold. First, to provide a biochemical framework for the activity of Rev1-Pol ζ at ICL lesions. While previous studies, including those from our own laboratories, have investigated the biochemical activities of Pol ζ and Rev1 at various lesions (9,30), these studies lacked the proper biochemical context, i. e. that of the 5-subunit Rev1-Pol ζ complex, together with its essential PCNA clamp, which may additionally by modified by ubiquitination at Lys-164. Secondly, we wanted to investigate which other DNA polymerase could participate in this process as a possible inserter enzyme, using both yeast Pol δ and Pol η as candidates. One firm conclusion from our studies is that, under all conditions tested, Pol δ was inhibitory to TLS, because of its tendency to degrade TLS insertion and extension intermediates with its proofreading exonuclease activity (Figure 4, 5B). In fact, as far as six nucleotides past the ICL, the lesion still affects the decision-making process by Pol δ, that of extension *versus* degradation (Figure 5B, lanes 14-16). We explored the possibility that ubiquitination of PCNA would reveal a unique step in TLS when multiple DNA polymerases were present. One of the proposed roles of ubiquitination is that it localizes Rev1, and by association Pol ζ to the lesion, perhaps favoring the competition between this TLS enzyme and the multitude of other PCNA client proteins for occupancy at the lesion. However, ubiquitinated PCNA did neither stimulate nor inhibit TLS by Rev1-Pol ζ in our studies, nor did it disfavor the inhibitory participation by Pol δ (Supplementary Figure S5). Studies of the role of PCNA ubiquitination in ICL-TLS in xenopus extracts were similarly uninformative (13).

From an enzymatic perspective, Pol η would be an ideal enzyme to serve as an inserter for ICL. In previous studies, Pol η has been shown to replicate past several model ICL lesions (33,49,50), and this is confirmed in the current study. Interestingly, in both the previous and our current studies, Pol η has the propensity to stall either after insertion at ICL, or just past the lesion (Fig. 1D). This would provide an ideal intermediate for a hand-off of intermediate from Pol η to Pol ζ or Rev1-Pol ζ. Our control studies show that, at least for this ICL substrate, the source of the inhibition could be attributed to an off-target reaction, i.e. the extension of the small ICL-associated covalent oligonucleotide by Pol η. While this Pol η-mediated extension of the ICL may also cause some inhibition of further bypass by Pol η itself (Figure 5A), more importantly, it forms an almost complete block for extension by Pol ζ or Rev1-Pol ζ (Figure 5B,C). Therefore, a two-enzyme TLS model involving Pol η and Rev1-Pol ζ would only be efficient if either the off-target extension were prevented or remedied, i.e. by a protein blocking extension of the unhooked ICL or a nucleases such as Pso2/SNM1A (51,52), specific for the attached ICL strand. Alternatively, the required strand displacement synthesis by Rev1-Pol ζ through the downstream double-stranded DNA could be facilitated for example by a DNA helicase. As a further complication, the genetic evidence for the involvement of Pol η (*RAD30*) in ICL repair in yeast is not strong. A yeast *rad30*Δ mutant showed no sensitivity to a variety of ICLs (14,53,54). Furthermore, loss of *RAD30* showed no effect on the hypermutability to cis-platin (55). By contrast, POLH deficient DT40 cells or XPV patients cells are mildly sensitive to cisplatin, consistent with a possible role of vertebrate Pol η in ICL repair, perhaps redundant with another polymerase (17,56).

Our studies with the five-subunit enzyme indicate that Rev1-Pol ζ may function as both an inserter and extender. Rev1 stimulated the synthesis of substantial percentage of extension products by Pol ζ, in a process which is only in part dependent on the catalytic activity of Rev 1 (Figure 2). Our data on the dispensability of Rev1’s catalytic activity are consistent with a genetic study, which shows that the catalytic inactive mutant of Rev1 does not sensitize yeast to treatment with cis-platin (48). Of importance for successful TLS is that the primer terminus is in close proximity to the ICL upon the recruitment of Rev1-Pol ζ (Figure 3). The function of Rev1 on non-damaged DNA is to inhibit DNA synthesis by Pol ζ, and this can be observed in all of our experiments. Therefore, engaging Rev1-Pol ζ too far from the ICL may result in premature stalling of the complex. Conversely, Rev1 also ensures that TLS is terminated soon after lesion bypass has been accomplished. While this type of regulation by Rev1 makes sense from a physiological point of view, our biochemical data cannot be easily be reconciled with a recent study, which suggests that Pol ζ can carry out extensive TLS in yeast, for stretches longer than 200 nucleotides, and introduce mutations during this process (57). Currently, it is not clear whether these extended mutagenic stretches are the result of processive DNA synthesis by Pol ζ, perhaps after dissociation of Rev1, or the result of iterative binding of Rev1-Pol ζ on the gapped, damaged DNA, resulting in close, but individual mutational patches.

Cells with mutations in Pol ζ are among the most hypersensitive to ICL-forming agents, and here we show that Rev1-Pol ζ can function as both an inserter and extender polymerase in bypassing ICLs. Our results further suggest that it is likely that in a physiological context, the insertion step across ICL is aided by another DNA polymerase such as Pol η. The limited sensitivity of Pol η deficient cells suggests that this function may be redundant with an additional TLS polymerase. The definitive answer to this question will have to await better biochemical systems to study ICL repair in yeast or human cells or conversely, improved genetics in *Xenopus laevis*, allowing for the generation of multiple simultaneous mutations in ICL repair genes in egg extracts that provide the only trackable biochemical system to study ICL repair to date. In the meantime, our studies provide a framework for the understanding of the role of Rev1-Pol ζ in ICL repair.

## Materials and Methods

### Reagents and Enzymes

Chemical reagents were purchased from Sigma-Aldrich (St. Louis, MO, USA). USER(tm) Mix, T4 polynucleotide kinase, and T4 DNA ligase were purchased from New England Biolabs (Ipswich, MA, USA).

### Oligonucleotides and Primers

The sequences of the oligonucleotides were listed in **Table S1**. HPLC-purified fluorescent labeled primers, biotinylated extension oligonucleotides, and splint oligonucleotides were purchased from Integrated DNA Technologies (Coralville, IA, USA). Single-stranded oligonucleotides with ICL precursors, 39mer 8 atom 20 bp ICL that contains two franking uracil bases around the crosslinked base (39+20(6) DMEDA ICL), and 39mer 8 atom 6 bp ICL (39+6 DMEDA ICL) were synthesized and purified as previously reported (33). The identity of the ICLs was confirmed by LC-MS. (**Figure S1A)**

### Preparation of 93 nt 6-bp ICL substrates

39+6 DMEDA ICL (500 pmol) was annealed with extension oligonucleotides (5’ext, 3’ext) and splint oligonucleotides (5’splint, 3’splint) in 1:1.2:2 ratio at room temperature overnight and reacted with T4 polynucleotide kinase (60 U) and T4 DNA ligase (4,800 U) at 37°C for 1 hour in the presence of 50 mM Tris-HCl (pH 7.5), 10 mM MgCl_2_, 1 mM ATP, 10 mM DTT. The product of the ligation reaction was purified by 10% denaturing polyacrylamide gel electrophoresis containing 7 M urea. The desired band was visualized by UV shadowing, excised and the DNA extracted by electroelution (Schleicher & Schuell BT 1000 or Bio-Rad Model 422). The extracted DNA was desalted by centrifugal filtration (Merck Millipore Amicon Ultra-0.5 mL), and freeze-dried. The purified product was analyzed by 10% denaturing polyacrylamide gel containing 7 M urea and 1X TBE. **(Figure S1B)**.

### Preparation of 93 nt 20-bp ICL substrates

A 93 nt-long single-stranded DNA that contains an ICL precursor (T93-C2) was prepared by ligating T39-C2 to 5’ext and 3’ext. The reaction and the purification methods were identical to those of 8a-6bp-ICL. The use of T39-G instead of T39-C2 gave the control undamaged 93mer substrate. An ICL was formed between T93-C2 and C20-C2 as previously (35). In brief, the two complementary oligonucleotides were annealed together, and then the ICL precursors were unmasked by sodium periodate to expose the aldehydes. The subsequent crosslink reaction by *N,N’-*dimethylethylenediamine (DMEDA) in the presence of sodium cyanoborohydride at pH 5.4 efforted the formation of the 8 atom ICL between T93-C2 and C20-C2. The purification method was identical to that 8a-6bp-ICL. The purified product was analyzed by 10% denaturing polyacrylamide gel containing 7 M urea and 1X TBE. **(Figure S1B)**

### Proteins

All proteins are the *Saccharomyces cerevisiae* species and were purified as previously described. Pol ζ, Rev1-Pol ζ, Rev1 and Rev1 mutants, Pol η, Pol δ and Pol δ-DV (D520V) were purified from yeast overexpression systems (5,6,36,58). RPA, PCNA and RFC were purified from *E. coli* overexpression systems (59-61).

### DNA polymerase assay

The 10 μl assay contained 40 mM Tris–HCl pH 7.8, 1 mM DTT, 0.2 mg/ml bovine serum albumin, 8 mM Mg-acetate, 100 mM NaCl final (this includes all contributions from the added enzymes), 0.5 mM ATP, DNA damage-response concentrations of dNTPs (36): 195 μM dCTP, 383 μM dTTP, 194 μM dATP and 50 μM dGTP, 10 nM control or ICL DNA substrate, 40 nM RPA, 30 nM PCNA, and 10 nM RFC. PCNA was loaded by RFC for 30 sec at 30 °C, and reactions were initiated with 25 nM of DNA polymerase. Assays containing the (mutant) Rev1-Pol ζ complex were initiated with 25nM Pol ζ and 75nM Rev1, which were preincubated on ice for 10 min or more. Aliquots were stopped with an equal volume of PK stop solution (20 mM EDTA, 1% SDS, 0.8 mg/ml Proteinase K), and incubated at 50° C for 30 min, followed by precipitation with an equal volume of 2.5 M NH_4_-acetate, 0.1 mg/ml glycogen. Then, 2.1 volumes of cold ethanol were added and, after cooling at −20 °C for 30 min, the DNA was collected at 17,000 x g in a refrigerated centrifuge for 20 min, followed by a 70% ethanol wash. The dried DNA pellet was re-suspended in 10 μl of 10 mM Tris-HCl pH 7.5, 1 mM EDTA, 50 mM NaCl, 0.2% SDS, and incubated at 50° C for 1 hour. An equal volume of sequencing solution was added to a final concentration of 50% formamide and10 mM EDTA, and the samples were analyzed on 12% polyacrylamide-7 M urea gels. Detection was carried out with a Typhoon phosphorimager in the fluorescence mode, and the data were quantified with ImageQuant (GE Healthcare), or with ImageJ software (after conversion of the Typhoon data by linearization). KaleidaGraph software was used for plotting. Note that the FAM-primers show variable ghosting of fluorescence at positions ∼1-2 nt higher. These ghosting phenomena are only for unreplicated primers and disappears upon replication.

### Polymerase exchange during TLS

The polymerase exchange assay was as the TLS assay, with modifications. The assay mixture contained 100μM AMP-CPP instead of ATP for PCNA loading by RFC (the α-β-methylene group allows proficient loading of PCNA but not replication by DNA polymerases when dNTPs are omitted) (62). Subsequently, DNA polymerase(s) were added to the assay, and, after 30 sec, replication was initiated by addition of dNTPs. Reactions were terminated after 15 min and processed as described above.

### Sequencing of the TLS synthesis products

Reactions were increased to 50 μl and allowed to proceed for 60min, stopped with final concentration of 50% formamide, 10 mM EDTA and 0.1% SDS, and loaded on a 12% polyacrylamide-7M urea preparative gel. Full-length products were extracted from the gel and purified by the ZR small RNA PAGE recovery kit (Zymo Research - R1070). Sequencing primer 5’-AAGCTGGAGCTCCACCGCGG-3’ was annealed and sequencing was performed with the USB Sequenase Version 2.0 DNA sequencing kit (Thermo Fisher Scientific - 70770).

## Acknowledgments

We thank Arnold Groehler IV (IBS Center for Genomic Integrity) for analyzing ICL-containing oligonucleotides by LC-MS. This work was funded in part by grants from the National Institutes of Health (GM118129 to PMB, CA165911 to ODS) and by the Korean Institute for Basic Science (IBS-R022-A1 to ODS).

## Author contributions

RBB, YKC, UR, ODS, and PMB planned this study. YKC and UR synthesized the substrates and RBB carried out the enzymatic studies. RBB, YKC, ODS, and PMB were involved in the interpretation of the results and the writing of the paper. All authors approved of the final version.

