## Supplemental data for "Bypass of DNA Interstrand crosslinks by a Rev1-DNA polymerase ζ complex"

[1] Department of Biochemistry and Molecular Biophysics, Washington University School of Medicine, Saint Louis, Missouri, USA, [2] Center for Genomic Integrity, Institute for Basic Science, Ulsan, 44919, Republic of Korea, [3] Department of Chemistry, Stony Brook University, Stony Book, NY 11794, USA, [4]Department of Biochemistry & Molecular Biophysics, Columbia University, New York, NY, 10032, USA, [5]Department of Biological Sciences, School of Life Sciences, Ulsan National Institute of Science and Technology, Ulsan, 44919, Republic of Korea

**Table S1. Sequences of DNA substrates and oligonucleotides**

| Name | Sequence |
|---|---|
| *DNA substrates:* | |
| Control undamaged 93mer | 5′-bio-CACTAGACGAAGCTTGATATGGGCGAAAGAAGGACAG<br>AAGAGGGTACCATCATAGAGTCAGTGGGCATTCAAGTGACGGGTACCATAGTCACG-3′-bio |
| 8a-6bp-ICL | 5′-bio-CACTAGACGAAGCTTGATATGGGCGAAAGAAG**G**ACAG<br>AAGAGGGTACCATCATAGAGTCAGTGGGCATTCAAGTGACGGGTACCATAGTCACG-3′-bio<br>,crosslinked to: 5′-pCT**G**TCC |
| 8a-20bp-ICL | 5′-bio-CACTAGACGAAGCTTGATATGGGCGAAAGAAG**G**ACAG<br>AAGAGGGTACCATCATAGAGTCAGTGGGCATTCAAGTGACGGGTACCATAGTCACG-3′-bio<br>,crosslinked to 5′-CCCTCTTCT**G**TCCTTCTTTC |
| *Fluorescent labeled primers:* | |
| P40-20 | 5′ FAM-CGTGACTATGGTACCCGTCACTTGAATGCCCACTGACTCT |
| P40 -8 | 5′ FAM-ACCCGTCACTTGAATGCCCACTGACTCTATGATGGTACCC |
| P40 -6 | 5′ FAM-CCGTCACTTGAATGCCCACTGACTCTATGATGGTACCCTC |
| P46 (0) | 5′ FAM-CCGTCACTTGAATGCCCACTGACTCTATGATGGTACCCTCTTCTGT**C** |
| P48 (+2) | 5′ FAM-CCGTCACTTGAATGCCCACTGACTCTATGATGGTACCCTCTTCTGT**C**CT |
| P52 (+6) | 5′ FAM-CCGTCACTTGAATGCCCACTGACTCTATGATGGTACCCTCTTCTGT**C**CTTCTT |
| *Biotinylated extension oligonucleotides:* | |
| 5′ext | 5′-bio-CACTAGACGAAGCTTGATATGGGC |
| 3′ext | 5′-phos-GGCATTCAAGTGACGGGTACCATAGTCACG-3′-bio |
| *Splint oligonucleotides:* | |
| 5′splint | 5′-TCTTTCGCCCATATCAAGCTTC |
| 3′splint | 5′-CACTTGAATGCCCACTGACTC |
| *39mer 8 atom ICLs:* | |
| 39+20(6) DMEDA ICL | 5′-GAAAGAAG**G**ACAGAAGAGGGTACCATCATAGAGTCAGTG, crosslinked to<br>5′-CCCTCTUCT**G**TCCUTCTTTC |
| 39+6 DMEDA ICL | 5′-GAAAGAAG**G**ACAGAAGAGGGTACCATCATAGAGTCAGTG, crosslinked to<br>5′-pCT**G**TCC-p3′ |
| *Template oligonucleotides:* | |
| T39-G | 5′-GAAAGAAGGACAGAAGAGGGTACCATCATAGAGTCAGTG |
| T39-C2 | 5′-GAAAGAAG**X**ACAGAAGAGGGTACCATCATAGAGTCAGTG |
| T93-C2 | 5′-bio-CACTAGACGAAGCTTGATATGGGCGAAAGAAG**X**ACAG<br>AAGAGGGTACCATCATAGAGTCAGTGGGCATTCAAGTGACGGGTACCATAGTCACG-3′-bio |
| C20-C2 | 5′-CCCTCTTCT**X**TCCTTCTTTC |

*Note: Bold, red G indicates the crosslink position of ICL; bold, red C: C opposite the crosslinked residue; bold **X**: ICL precursor; bio: biotin; p: phosphate; FAM: fluorescein.

**A.**

| Name | Calculated Mass | Observed Mass | Accuracy |
|---|---|---|---|
| 39+20(6) DMEDA ICL | 18209.175 | 18209.1655 | ±0.522 ppm |
| 39+6 DMEDA ICL | 14230.497 | 14230.4928 | ±0.295 ppm |

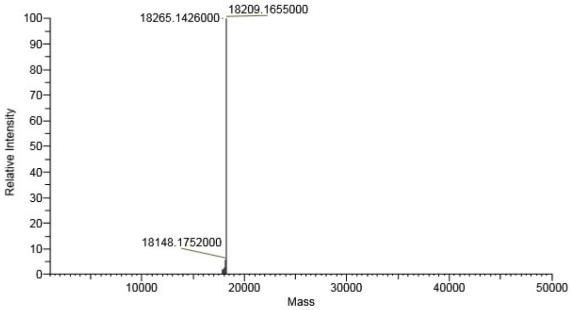


Deconvoluted spectrum of
39+20(6) DMEDA ICL

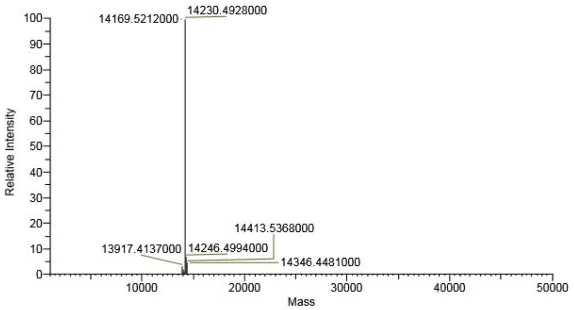


Deconvoluted spectrum of
39+6 DMEDA ICL

**B.**

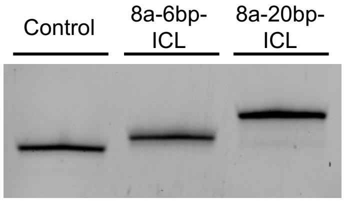


**Figure S1. Analysis of the DNA substrates used in this study. (A)** LC-MS analysis of ICL intermediates. UPLC-ESI$^-$-MS analyses were conducted with a Thermo Scientific Q-Exactive Focus hybrid quadrupole-Orbitrap mass spectrometer interfaced with a Thermo Scientific UltiMate 3000 UHPLC system. A heated electrospray ionization (HESI-II) probe was operated in negative ionization mode. The analyzer was operated in full-MS mode with a scan range of *m/z* 200 – 2000. A Thermo Scientific Hypersil GOLD (100 x 2.1 mm, 1.9 µm) column was conditioned with 15mM ammonium acetate and acetonitrile at 40 °C at a flow rate of 50 µL/min. DNA oligo (50 pmol) was eluted with a gradient that was initially kept at 2% acetonitrile for 2 minutes, then linearly increased to 80% acetonitrile over 16 minutes, kept at 80% acetonitrile for 2 minutes, decreased to 2% acetonitrile over 2 minutes, and finally re-equilibrated at 2% acetonitrile for 13 minutes. Under these conditions, DNA oligos eluted between 14 – 16 minutes. Spectral data was deconvoluted using Thermo Scientific BioPharma Finder 3.1 software. **(B)** 0.5 pmol of the purified 93mer substrates (Figure 1A, Table S1) were loaded to a 10% polyacrylamide gel containing 7 M urea and 1X TBE. The resolved gel was stained with SYBR gold (Thermo Scientific) and visualized by biomolecular imager (GE Typhoon FLA 7000).

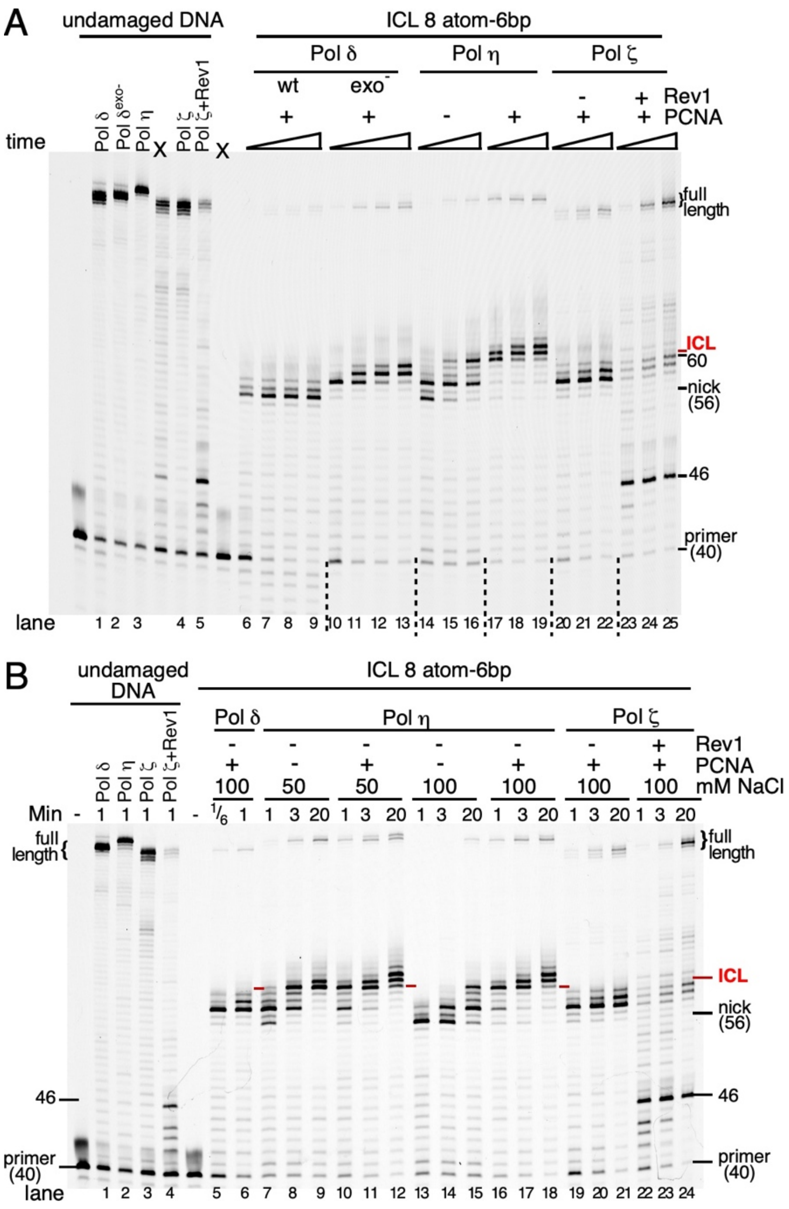


**Figure S2. Translesion DNA synthesis of an ICL by yeast DNA polymerases. (A)** This is the entire gel, which was cut up for easier visualization in Figure 1C. Lanes removed are indicate with an X. **(B)** Time course of replication of control DNA (left section) or ICL DNA. Assays with Pol η were performed at the indicated concentrations of NaCl and either with or without PCNA. Assays with Pol δ and Pol ζ, the latter with or without Rev1, were performed at 100 mM NaCl.

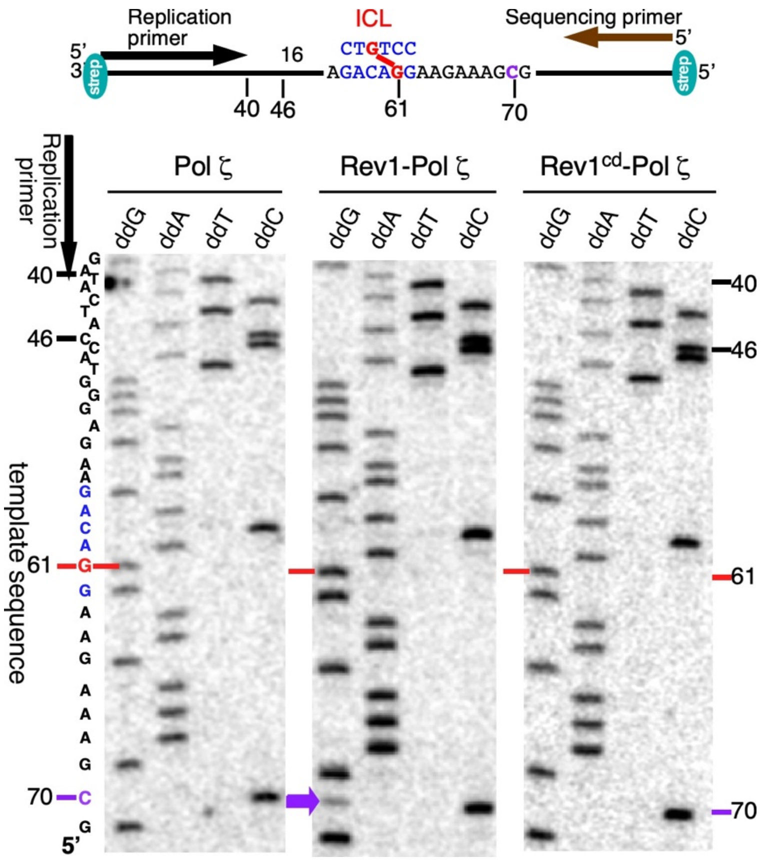


**Figure S3. Sequencing of TLS products.** Extended sequencing reaction of the section shown in Fig. 2C. Top, schematic map of the assay. Full-length replication products were isolated from 60 min assays. Bottom, sequencing analysis. The red arrow indicates the ICL-dG template position, which is replicated by insertion of dCMP, which is sequenced as a dG. The purple arrow at position 60 indicates a position where Rev1, but not Rev1$^{cd}$ misincorporates dCMP across from template dC with a frequency of 15-20%. Note that at the major template dC stall site at position 46, dGMP is faithfully incorporated opposite the dC template.

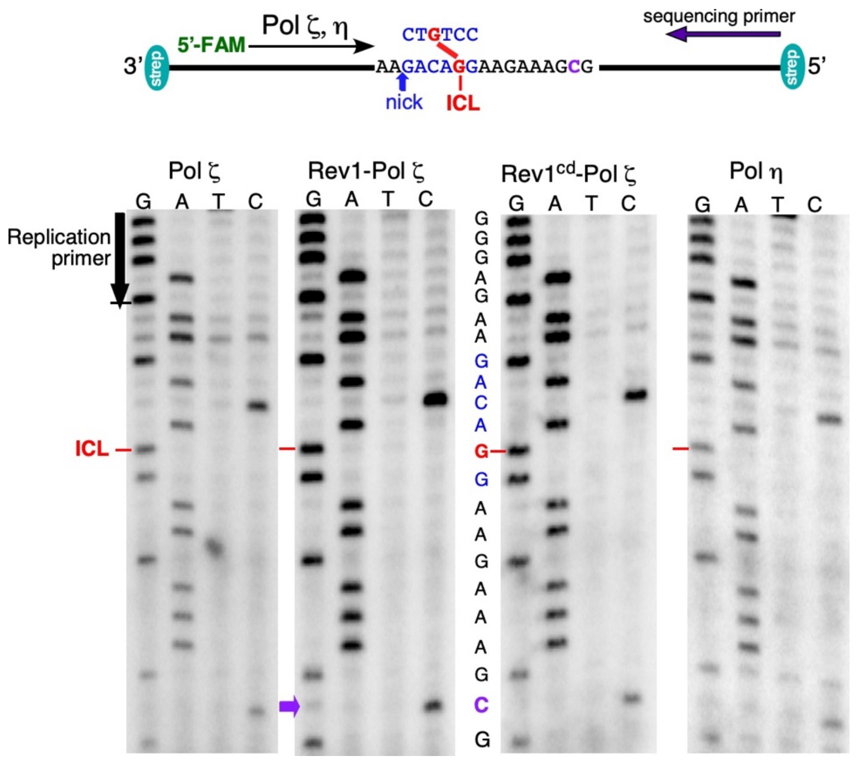


**Figure S4. Sequencing of TLS products.** Top, schematic map of the assay. See Fig. 3 for TLS assay. Bottom, sequencing analysis. Full-length replication products were isolated from 60 min assays. The red arrow indicates the ICL-dG template position, which is replicated by insertion of dCMP, which is sequenced as a dG. The purple arrow at position 60 indicates a position where Rev1, but not Rev1$^{cd}$ misincorporates dCMP across from template dC with a frequency of 15-20%.

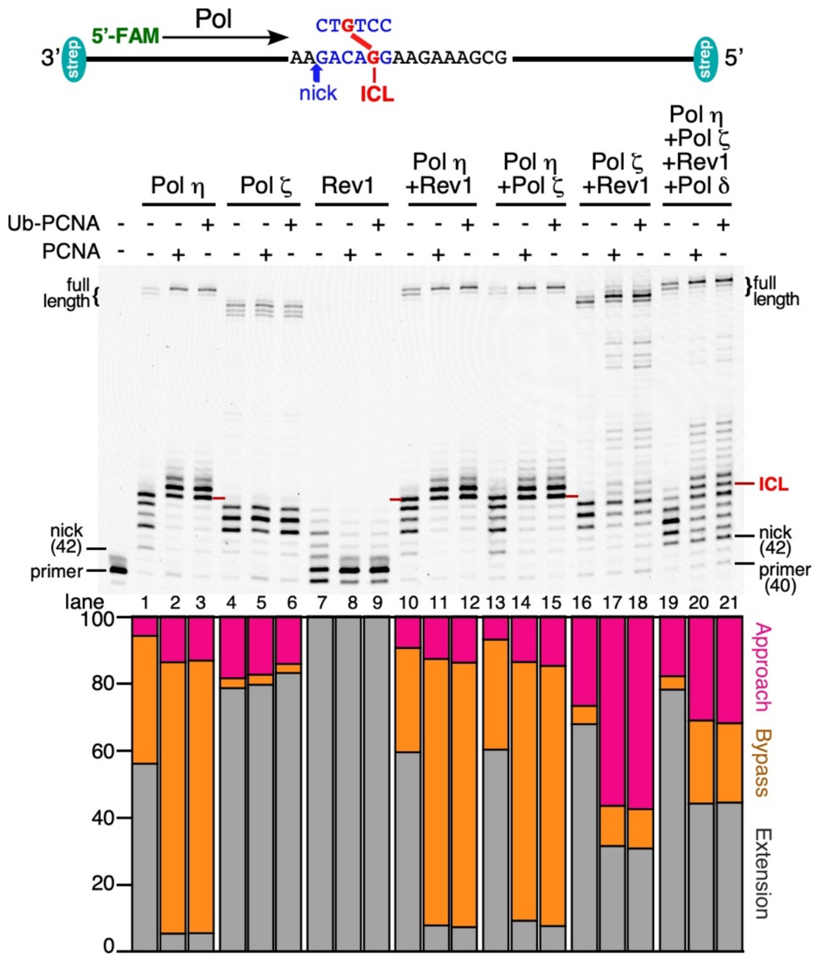


**Figure S5. Stimulation of ICL TLS by PCNA versus ubiquitinated PCNA.** Top, description of the substrate. Middle, the assay is as described in Figure 4, with either no PCNA, or 30 nM PCNA or ubiquitinated PCNA (as trimer), for 15 min at 30 ˚C. bottom, quantification of products in three classes.
